# Protein Pocket Coengagement Prioritizes Drug–Drug Interactions

**DOI:** 10.64898/2026.10.28.762582

**Authors:** Jeffrey Skolnick, Bharath Srinivasan

## Abstract

Drug–drug interactions (DDIs) are largely catalogued empirically, with limited structural explanation of why two drugs interact. We asked whether a purely structural mechanism — competing or cooperating engagement of the same protein pocket — is sufficient to recover documented DDIs without any training on clinical interaction labels. Using composite protein multiple-ligand (COLIG) assignments, which classify co-bound ligand pairs as attractive or repulsive, we scored 21,945 pairs of 210 drugs by predicted engagement of each protein’s top-ranked pocket. Confidence-summed COLIG ranking achieved top-5% precision of 49.3% against DDInter and 25.1% against KEGG, versus background prevalences of 29.6% and 14.7%, respectively; pooled top-pocket engagement reached 50.3% DDInter precision. These results support pocket-based DDI screening and class-dependent hypotheses about direct and noncontact ligand engagement, while leaving causal mechanisms to be established.

## Introduction

A drug–drug interaction (DDI) occurs when coadministration changes the disposition or effect of one or both drugs. Outcomes can include altered exposure, enhanced toxicity, reduced efficacy, or potentiation of an intended response. Pharmacokinetic interactions affect absorption, distribution, metabolism, or excretion and thereby change the concentration of a drug or its active metabolites. For example, inhibition of a drug-metabolizing enzyme can increase substrate exposure, whereas induction can reduce it; effects on prodrug activation can differ.^1,2^ Pharmacodynamic interactions alter the combined response through shared targets or coupled physiological processes, potentially enhancing a response, opposing it, or combining toxicities. A clinical potentiation annotation does not by itself establish synergy under a quantitative combination-response model.^3,4,5^ Identifying a documented DDI is therefore different from predicting its severity, physiological consequence, or therapeutic synergy.

Clinical interaction resources provide practical benchmarks for identifying documented DDIs. DDInter integrates published evidence and drug-label information with severity and mechanism annotations, and DDInter 2.0 expands its coverage and interaction types.^6,7^ The Kyoto Encyclopedia of Genes and Genomes (KEGG) also provides interaction information; its interaction queries retrieve contraindication and precaution relationships from Japanese drug labeling.^8,9^ These resources summarize heterogeneous clinical and pharmacological evidence rather than a uniform assay of direct drug–drug contact. We therefore treat database nonlisting as an unreported interaction, that is not necessarily an experimentally established negative.

Several approaches infer DDIs by transferring information between related drugs. Vilar and colleagues combined molecular fingerprints with known interaction records to nominate interactions involving chemically similar compounds.^10^ INDI integrated chemical and biological similarities to infer pharmacokinetic and pharmacodynamic interactions and associated recommendations.^11^ Later approaches used machine learning to predict interaction labels or types from drug representations.^12,13^ These studies established the value of chemical and biological representations, but their dependence on existing interaction or biological annotations differs from a fixed ranking constructed from predicted protein engagement without fitting DDI labels. Published performance estimates are consequently not directly comparable with the present benchmark.

Protein annotation resources connect structural assignments to biological targets. UniProtKB supplies sequences, stable accessions, protein names, and functional annotations; its reviewed Swiss-Prot records provide the human protein mapping used here.^14^ SIFTS (Structure Integration with Function, Taxonomy and Sequences), maintained by the Protein Data Bank in Europe and UniProt teams, links structural chains and residues to sequence entries.^15^ These resources identify and annotate proteins; they do not supply the clinical DDI labels used for evaluation.

Target relationships provide another route from molecular properties to drug effects. Keiser and colleagues related proteins through the chemical similarity of their ligands and experimentally confirmed unexpected pharmacological associations.^16^Lounkine and colleagues subsequently tested predicted activities of marketed drugs on unintended targets, demonstrating extensive polypharmacology and a route to investigating target-associated adverse effects.^17^ These studies support examining the ensemble of proteins engaged by a drug rather than assuming a single therapeutic target. They do not imply that two drugs sharing a target must interact clinically.

Drug pairs may also influence different proteins that are functionally coupled. Huang and colleagues found shared-target enrichment and an association between pharmacodynamic DDIs and protein-network proximity using known DrugBank target relationships.^18^ Decagon used a graph convolutional model to predict specific polypharmacy side effects from protein interactions, drug– target links, and drug-pair side-effect records.^19^ Such studies establish the usefulness of network representations under their respective input conditions and endpoints. Whether an explicit network adds information beyond a structural engagement score requires a matched comparison.

Pocket structure permits a more local description of a candidate interaction. A composite protein multiple-ligand interaction, termed a COLIG, describes ligands that interact with one another while occupying the same protein pocket. Tonddast-Navaei and colleagues combined structural analysis with experiments on dihydrofolate reductase to demonstrate the relevance of such interactions.^20^ Pocket-comparison methods such as APoc and structural resources such as BioLiP2 provide complementary foundations for examining ligand engagement across proteins.^21,22^ Here, attractive and repulsive predictions of the pair of ligands present in a given COLIG are combined as unsigned COLIG evidence. Their structural sign is not a clinical label: attractive does not mean beneficial, and repulsive does not mean harmful or pharmacodynamically antagonistic.

We distinguish COLIG evidence from NoCOLIG assignments, in which both drugs are assigned to the same pocket without a predicted direct ligand–ligand interaction. Such engagement could still influence protein function. The Monod–Wyman–Changeux model provides a physical framework in which ligand binding shifts protein conformational states and mediates allosteric coupling, but it does not establish that our NoCOLIG assignments are allosteric.^23^ Direct ligand–ligand interactions and noncontact pocket engagement are therefore plausible, potentially coexisting routes to a functional response.^20^ Comparing their DDI enrichment across protein classes can guide experiments without assuming that a predicted assignment establishes occupancy, inhibition, or activation.

Metabolites provide another source of distributed regulation. Proteome-scale mapping by Piazza and colleagues demonstrated extensive protein–metabolite interactions beyond conventional enzyme–substrate assignments.^24^ The entabolon framework proposes that shared metabolite binding couples proteins and contributes to collective pathway behavior.^25^ A drug-pair score aggregating pocket evidence across distinct proteins may therefore summarize distributed engagement even without explicit signal propagation. STRING and metabolite-coupling networks provide complementary representations for testing whether broader connections improve prediction.^25,26^ Their relevance here is an empirical question; a structural or pathway assignment does not demonstrate perturbation in a particular tissue.

Clinical pharmacokinetic interactions illustrate how an effect at an enzyme or transporter can alter whole-body drug exposure. In a randomized crossover study, ketoconazole and itraconazole increased midazolam exposure, consistent with inhibition of CYP3A-mediated metabolism.^2^ Clinical digoxin interactions have also been compared with in vitro P-glycoprotein inhibition, and rifampicin inhibits organic-anion uptake in experimental human liver transport systems.^27,28^ These examples support local molecular engagement as a potential initiating event, but do not demonstrate simultaneous pocket occupancy or contact in a predicted COLIG geometry. Our structural assignments nominate candidate relationships rather than establish the mechanism of an individual clinical interaction.^1^

Here, we evaluate fixed pocket-based rankings over all pairs in a defined drug cohort, emphasizing precision among the top-ranked 5%. A central comparison distinguishes attractive or repulsive COLIG assignments from explicit same-pocket NoCOLIG controls. We report protein-class subsets alongside complete-cohort performance, separate distinct supporting-protein count from mean confidence, and test network and pathway additions. The primary rankings do not fit clinical interaction labels; secondary supervised combinations use drug-disjoint evaluation. These analyses assess preliminary interaction screening and class-dependent functional hypotheses, leaving physiological consequences and causal mechanisms for subsequent validation.

## Materials and methods

Figure 1 summarizes the analysis. LIGMAP predicts small-molecule engagement of protein pockets, and the contact scheme described below classifies predicted ligand–ligand relationships as attractive or repulsive.^20,25^ Evidence for each drug pair is aggregated across distinct proteins and examined in the context of protein folds, interaction networks, and pathways. Fixed scoring functions rank the drug pairs, and performance is assessed primarily by the fraction of the top-ranked 5% listed as interacting in each reference database.

**Figure 1.**
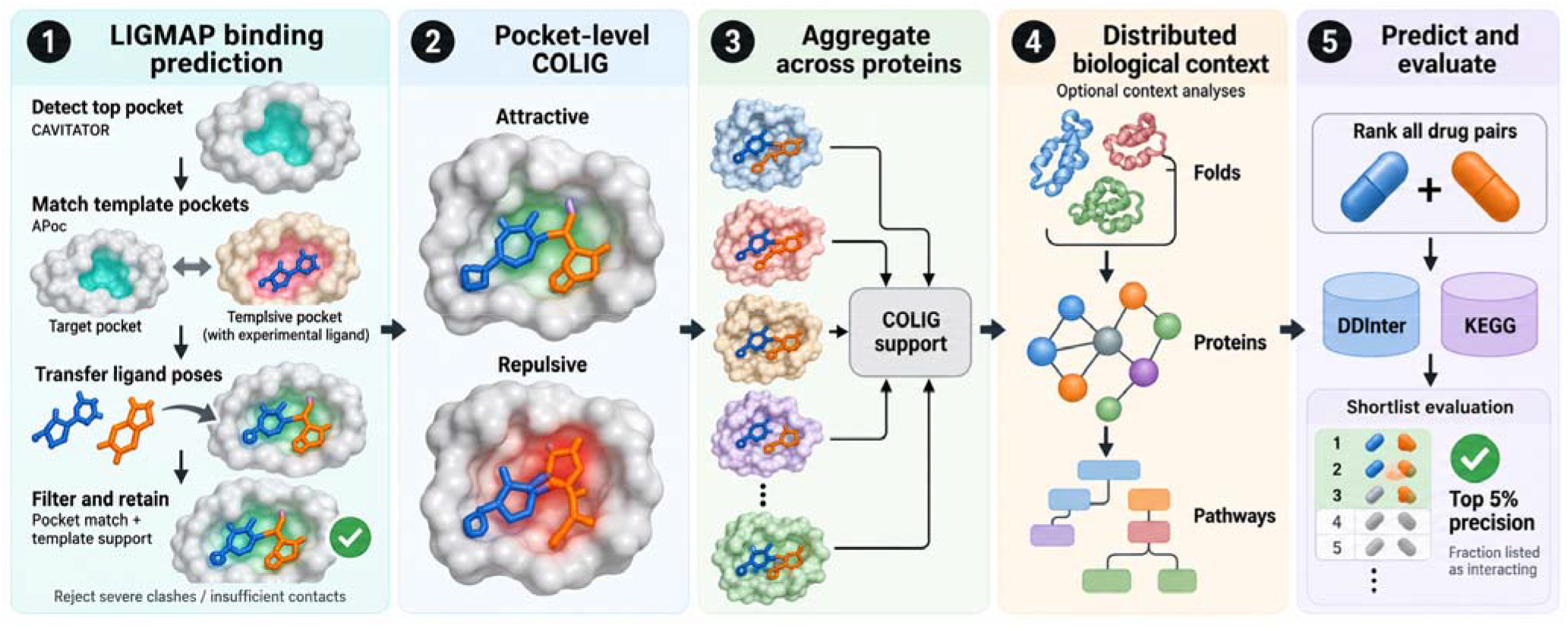
Overview of structural COLIG screening. LIGMAP transfers template-ligand information to target pockets using pocket sequence and structural similarity. Attractive and repulsive ligand–ligand assignments are aggregated across distinct proteins to rank drug pairs for comparison with DDI references. Network and pathway analyses provide optional biological context; they are not required components of the fixed local score. Folds denotes protein folds.

LIGMAP predicts small-molecule binding by transferring information from experimentally determined protein–ligand complexes to structurally similar target pockets. CAVITATOR identifies the target’s largest pocket, and APoc compares it with a library of ligand-bound template pockets.^21,25^ Candidate ligands are selected using pocket-similarity significance, conservation of aligned residues, and support from multiple templates. Pocket alignment supplies the transformation used to place each template ligand in the target structure. Geometric filters reject placements with excessive protein overlap or insufficient protein contact. Predicted ligand placements then provide the basis for classifying putative COLIG contacts. The simplified, symmetric atom-type scheme treats carbon– carbon and oxygen–nitrogen contacts as attractive and carbon–oxygen, carbon–nitrogen, nitrogen– nitrogen, and oxygen–oxygen contacts as repulsive. Sulfur is assigned the oxygen interaction type, and phosphorus the nitrogen type. These classifications are qualitative contact scores, not quantitative interaction energies.

### Study design and prediction releases

The cohort comprised 210 mapped drugs (Table S1) and every unordered pair of different drugs, giving N = 210 × 209/2 = 21,945 pairs. No pair was removed because its interaction status was unknown or its structural score was zero. The primary fixed scores were constructed without fitting DDI labels, making the main analysis a retrospective evaluation of a fixed ranking rather than a supervised train–test experiment. Repeated exploratory examination of this benchmark does not substitute for prospective validation.

### Clinical interaction references

The frozen DDInter reference contained 6,495 listed interactions and the KEGG reference contained 3,219, corresponding to 29.5967% and 14.6685% of all candidate pairs, respectively (Tables S2 and S3). A positive label means that a pair is recorded in the evaluated reference, not that it passed a uniform assay of direct molecular interaction. DDInter curates literature and drug-label information, while the KEGG interaction operation extracts contraindication or precaution relationships from Japanese labeling.^6,7,9^ Neither provides a complete experimentally established negative set. Unlisted pairs were therefore designated unreported rather than true negatives. Interaction severity and anticancer synergy were not the study endpoints.

A saved DDInter mechanism/API snapshot contained 8,555 listed pairs in the same cohort, including all 6,495 frozen-reference positives and 2,060 additional pairs (Table S4). We call this the expanded DDInter reference. Its greater coverage reflects database availability, not newly discovered interactions or experimental confirmation by this analysis. KEGG provides a second reference representation, but shared source material prevents treating the two resources as fully independent clinical evidence.

### Structural records and protein mapping

Attractive and repulsive inner archives list ligand pairs, whereas combined archives list individual ligand predictions; archive headers declare record counts. Ligand Chemical Component Dictionary identifiers were mapped to the frozen 210-drug dictionary, and self-pairs and unmapped identities were excluded without expanding the cohort. The Protein Data Bank (PDB) supplies experimentally determined macromolecular structures and archived SIFTS chain-to-UniProt mappings connect those structures to sequence accessions.^15,29^ Reviewed UniProtKB/Swiss-Prot annotations supplied human protein symbols, families, enzyme classifications, and controlled keywords.^14^ AlphaFold 2 identifiers were joined by UniProt accession.^30^ Repeated structures, contexts, and isoforms mapped to the same protein were deduplicated for the primary score. When a multichain record mapped to multiple human proteins and the ligand-bearing chain was not retained, support was provisionally assigned to the mapped proteins; this archive ambiguity limits interpretation. The current release contained 131,051 unordered drug-pair–protein supports and 3,916 proteins in the union of predicted binding and COLIG records. Structure–sequence correspondence and protein annotation do not establish ligand occupancy or a DDI.

The archived rows contain predicted relationships and supplied confidence values. They do not constitute measured affinities, ligand concentrations, or occupancy probabilities.

### Scoring drug pairs from distinct supporting proteins

For each drug pair a,b and supporting protein g, the unsigned confidence q retains the maximum supplied value across attractive and repulsive records and structural contexts (eq 1). The local score sums these retained confidences across distinct proteins, whereas the count score records the number of proteins with positive support (eq 2).

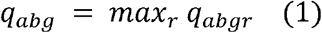

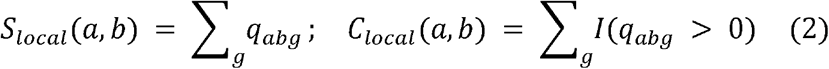

We distinguish three types of evidence: attractive or repulsive COLIG assignments within the top-ranked pocket; explicit NoCOLIG assignments to that pocket without a predicted direct ligand– ligand interaction; and pooled top-pocket engagement, which ignores the interaction classification. None of these scores is a calibrated probability of clinical interaction. Evaluation measures recovery of reference-listed DDIs. Attractive and repulsive protein counts are retained separately, and their supporting sets can overlap. A zero score indicates absence of support in the supplied prediction records, not evidence that the drugs do or do not interact.

Binding-burden features use the smaller and larger target-set sizes, shared-target count, and Jaccard similarity. In this setting, target overlap retains the top-pocket restriction of its input records and is not a pocket-independent control. However, the saved overlap score intersects target sets after aggregation to proteins. Shared protein membership alone does not independently verify a common structural record or identical pocket geometry across different structures. We therefore describe this as a protein-level proxy for pooled top-pocket engagement, reserving the explicit COLIG versus NoCOLIG comparison for structure-matched assignment types. In the same-pocket NoCOLIG analysis, unordered drug pairs from corresponding singleton pocket lists were retained only when absent from attractive and repulsive records for that same structural context. A pair’s COLIG on another structure did not invalidate the observation. The pooled comparator must not be interpreted as NoCOLIG-only evidence.

### Ranking metrics and random backgrounds

Let U denote the evaluated drug-pair population, s□ a fixed structural score, and y□ the indicator that pair i is listed in the evaluated reference. We rank scores from largest to smallest and select the top 5% of the eligible pair population. In the full cohort, results in 1,097.25 pairs. If the cutoff falls within a tied group, every pair in that group receives the same fractional weight so that the total selected weight equals the budget exactly. Precision is the fraction of selected pair weight carrying a reference-positive label; recall is the fraction of all reference-positive pair weight recovered. Background precision is the reference-positive fraction of the evaluated population, and enrichment is precision divided by that background. Thus, 1.67-fold enrichment means 1.67 times the reference-positive fraction expected under uniform pair selection. It is neither a P value nor a probability of causation. Precision–recall curves sweep tied-score groups without using labels to reorder ties.^31^

Uniform-pair sampling supplies a descriptive chance reference. For a support set of n selected pairs with k reference positives, the upper hypergeometric tail conditions on the eligible universe and its number of positives. Mechanism analyses also used 20,000 hypergeometric draws preserving fractional threshold ties. These calculations assume exchangeable pairs and do not preserve drug-specific annotation rates; dependence-aware intervals are therefore reported alongside them. Scores with fewer supported pairs than the screening budget require zero-score padding and are flagged rather than presented as fully supported screens.

### Chemical group resampling and supervised comparisons

We grouped drugs using binary radius-2, 2,048-bit Morgan fingerprints and Tanimoto similarity, T = c/(a + b − c), where a and b count set bits and c counts shared bits. Connected groups used a similarity threshold of 0.8, giving 209 groups for 210 drugs. Chemical structure defined resampling and split groups, not the primary ranking.^32,33^

For 1,000 multinomial resamples of whole chemical groups, each pair received the product of its endpoint-group multiplicities. Scores were reranked within the applicable resampled population, using the same draw for paired comparisons. The 2.5th and 97.5th percentiles form the reported 95% intervals. These intervals are conditional on the available predictions and exclude upstream prediction, mapping, and model-refitting uncertainty. The legacy analysis used 5,000 resamples and is identified separately.

Secondary supervised comparisons used three fixed repetitions of five-fold chemical-group partitions. Each unordered pair of test folds defined one test block; training excluded every pair containing a drug from either test fold. This produced 15 blocks per repetition, 45 fits per model and endpoint, and three held-out predictions per evaluated pair. Both drugs and their chemical groups were absent from the corresponding training data. Scaling and fitting used training data only.

The current logistic baseline used log-transformed attractive and repulsive protein counts, target-burden and overlap features, and attractive/repulsive drug–metabolite similarities. Network and pathway additions used the same partitions. Logistic regression used L2 regularization, C = 0.1, and at most 2,500 iterations; mechanism-specific local models additionally included unsigned protein count and confidence sum. Additional nonlinear and count-only comparisons are identified in the supplementary results. No claim of comprehensive algorithm optimization is made.

The drug–metabolite deliveries contain 31,994 deduplicated context/sign/drug/metabolite/protein observations derived from the LIGMAP predictions. Per-drug profiles retain the maximum confidence per metabolite or protein–metabolite combination, depending on the analysis. Attractive and repulsive profiles are separate. Metabolite-identity similarity uses inverse-frequency weighting and, in the confidence variant, the supplied profile weights. Distinct-metabolite count counts each identity once across proteins and structural contexts. These features describe predicted structural engagement, not measured metabolite depletion or pathway activity.

STRING integrates experimental and curated evidence for protein associations; the present calculation uses its physical-interaction layer.^26^ Reactome supplies literature-curated reaction and pathway membership, not measured pathway activity in treated cells.^34^ These biological annotation layers are distinct from the clinical DDI labels.

### Consideration of mechanisms of drug-drug interactions

DDInter mechanism flags define pharmacokinetic interactions as the union of absorption, distribution, metabolism, and excretion; potentiation uses the database synergy flag, and antagonism uses its antagonism flag. Unspecified mechanism denotes records lacking all six flags, including records marked other. Flags can overlap. Each main mechanism-specific population contains frozen-reference positives for that category plus the 13,390 pairs unlisted in both DDInter snapshots. Other known interactions are excluded rather than used as presumed negatives for the category. Expanded-positive, legacy-background, full-universe retrieval, and within-known-DDI sensitivities are reported separately. Five models for each of four main categories, evaluated over 45 held-out blocks per model, required 900 fits. A clinical potentiation annotation is not equivalent to synergy measured against Bliss independence or Loewe additivity.^3,4,5^

The FDA clinical examples independently identified 28 P450-substrate drugs, defining 5,474 pairs containing at least one such drug.^35^ Inhibitor and inducer assignments were retained separately and were not treated as evidence of substrate status. This example list is not exhaustive. The core P450 panel comprised CYP1A2, CYP2A6, CYP2B6, CYP2C8, CYP2C9, CYP2C19, CYP2D6, CYP2E1, CYP3A4, and CYP3A5. FDA CYP3A annotations were interpreted at the family level rather than as proof of a particular isoform. Strict enzyme alignment required structural support on a documented substrate enzyme for at least one partner. Shared drug–metabolite support required matching protein and metabolite identities.

### Questions addressed by the comparisons

The comparisons address three related questions, with clinical labels used only for evaluation of fixed rankings or for training within the explicitly identified supervised analyses.

1. Screening enrichment: does the highest-scoring 5% contain a larger fraction of documented DDIs than uniform selection from the same eligible population? Top-5% precision is compared with the reference prevalence in that population.
2. Structural specificity: do attractive or repulsive COLIG assignments add information beyond assigning both drugs to the same predicted top pocket? Explicit NoCOLIG controls lack a predicted direct ligand–ligand interaction in the matched structural unit.
3. Additional biological context: do metabolite coupling, STRING physical interactions, or their combination improve ranking beyond local pocket evidence? Fixed propagation and rank-fusion scores do not fit DDI labels; secondary supervised combinations use the drug-disjoint evaluation described above.^25,26^

### Explicit same pocket controls across protein classes

We extended the kinase comparison to all 21 primary protein target classes represented in the supplied predictions. These are protein classes, not therapeutic drug classes. Attractive and repulsive pair records were combined as unsigned COLIG evidence. Explicit NoCOLIG archives supplied individual drugs assigned to the same pocket without a predicted direct ligand–ligand interaction, according to the protein rating assignment definitions. Unordered pairs of distinct mapped drugs were formed within each file. A NoCOLIG pair was excluded if it occurred in attractive or repulsive records for that same structural context and canonical structure identifier. A COLIG on another structure or protein did not invalidate the local NoCOLIG observation. The archive assignment unit was used without inferring an undocumented additional pocket identifier or independently verifying poses.

For each assignment type, repeated records were collapsed by maximum confidence per pair and protein. COLIG confidence used the supplied pair value; NoCOLIG confidence used the smaller of the two supplied singleton values before the same maximum-per-protein and sum-across-proteins aggregation. This minimum is a scoring proxy, not a calibrated joint binding probability. Distinct supporting-protein counts supplied a sensitivity analysis.

Each class comparison used every drug pair sharing at least one predicted target in that class, based on the combined single-drug records, and its own reference prevalence. This is a narrower population than the both-drugs-bind-class populations of Table 5. Every observed class COLIG and NoCOLIG pair fell within its corresponding shared-target population. Drug pairs could occur in several classes even though primary protein classes were mutually exclusive. The all-protein summary pooled unique proteins and pairs; it was not an average of class precisions.

We retained the top-5% precision where both assignment types had enough support to fill the budget. Equal top-100 comparisons required at least 100 supported pairs for both types. A descriptive smaller-budget analysis used K = min(100, number of COLIG-supported pairs, number of NoCOLIG-supported pairs), restricted to K at least 20. Eligibility was determined from coverage without clinical labels and does not imply adequate statistical power. Fractional boundary ties and the same 1,000 chemical-group bootstrap resamples were used. Bootstrap samples could have insufficient control mass even when the observed data filled the budget; these padding frequencies were retained.

Paired percentile intervals were supplemented by simultaneous intervals from the maximum absolute standardized centered bootstrap error across eligible primary classes, separately by endpoint, score, and budget. Overlapping all-protein and enzyme summary sets were excluded from that classwise adjustment. This adjustment does not cover all prior exploratory analyses. Equal budgets do not eliminate target-burden, confidence-calibration, or clinical-ascertainment differences.

The mean confidence score measures the average retained structural confidence per distinct supporting protein for a drug pair. It separates the average strength of the protein-level evidence from the number of proteins providing that evidence:

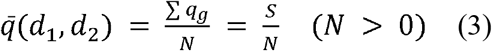

For drug pair *d1, d2, q*_*g*_ denotes the retained confidence for supporting protein g, N is the number of distinct supporting proteins, and S is the sum of the retained protein-level confidences.

When several structures or structural contexts support the same drug-pair-protein combination, the analysis retains only the maximum confidence for that protein before calculating S and N.

### Matched structural units and molecular association contexts

We distinguished four structural contexts: isolated PDB monomers, AlphaFold2 monomers, noninterface single-chain pockets within dimers, and multimer/interface-adjacent dimer pockets. Only the top-ranked pocket was reported, and the delivered assignment lists do not retain coordinates or additional pocket identifiers. COLIG denotes a predicted ligand–ligand interaction within that pocket; NoCOLIG denotes same-pocket assignments without that predicted interaction. Assignments may differ for the same drug pair across proteins or structures. The pooled structural-unit population is stricter than the protein-intersection proxy used in the class-conditioned analysis.

To isolate assignment type from differences in confidence definitions, we assigned every paired unit the same common weight, min(q_a,q_b), from its combined singleton records. Pooled, COLIG and NoCOLIG scores all used the maximum weight per pair/protein and then summed over distinct proteins. Counts used the same retained pair/protein supports. Native supplied COLIG pair confidence and native NoCOLIG singleton minima were retained as separate sensitivities. Neither singleton minima nor the supplied scores are independently calibrated joint occupancy probabilities. We evaluated all scores over the full 21,945 drug-pair universe, the identical matched pooled population, and equal numbers of min(100, COLIG-supported pairs, NoCOLIG-supported pairs), requiring at least 20 pairs for the latter. Zero-score padding was flagged rather than interpreted as supported prediction.

We repeated fixed rankings by context, after excluding each context, and after excluding multiprotein structural units. A second coverage sensitivity compared isolated PDB monomers with non-interface dimer chains using only proteins represented in both. We allocated each selected pair/protein maximum to its supporting context, dividing exact ties equally. This decomposition describes score composition rather than causal importance. Paired uncertainty used 1,000 chemical-group bootstrap draws with seed 20261018, recomputing the weighted rankings. These additional context and class comparisons are exploratory and use pointwise intervals.

### Jackknife and cross resource validation sensitivities

We removed each drug in turn and repeated the fixed rankings. A second jackknife removed all members of each of the 209 chemical groups in turn; neither analysis refitted score parameters. We also compared rankings with saved Liverpool COVID-19, HIV, and hepatitis interaction assessments.^36,37,38^ Drugs were matched by exact normalized names without inferring ambiguous or combination mappings. Red, amber, or yellow warnings were contrasted with explicit green assessments, with a second endpoint excluding yellow. Contradictory green/risk records were excluded within each source and from the combined analysis. Unlisted pairs were not considered. These expert assessments can reflect pharmacological expectations rather than uniformly measured interaction or noninteraction outcomes.

### Network propagation and metabolite sparsification

To examine whether explicit cellular-network context improves DDI ranking, we constructed metabolite-coupling and STRING protein-interaction graphs, propagated drug-target confidence through them, and ranked pairs by the similarity of their propagated profiles. Fixed-score combinations did not fit clinical DDI labels.

### Metabolite and STRING transition matrices

Let *p*_*gf*_ denote the supplied confidence linking protein g to metabolite feature f and let *d*_*f*_ be the number of proteins associated with that feature. A feature represents a single metabolite or a metabolite pair. For distinct proteins, the metabolite-protein adjacency matrix is

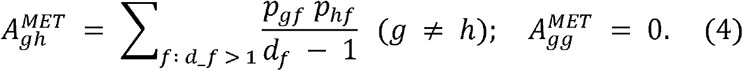

Self-edges are removed. We defined the row normalization by dividing each entry by its row sum when that sum is positive:

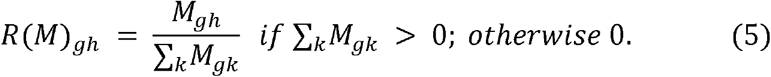

STRING version 12 physical links with scores of at least 700 define a separate adjacency matrix *A*^*STR*^. The metabolite and STRING matrices were normalized separately, and we then row-normalized their sum for combined network propagation: ^26^

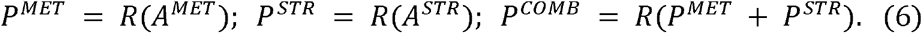

### Drug seeding and propagation

For drug a, let *C*_*ga*_ denote its predicted target confidence on protein g. For drugs with positive total seed confidence, normalize the seed vector to unit mass:

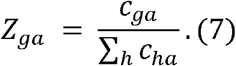

Let Z collect these drug seed vectors as columns. Starting from *X* ^(0)^ = Z, apply 40 updates using the selected transition matrix P:

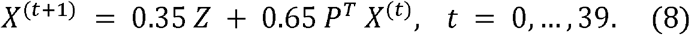

The fixed network score is the cosine similarity between the resulting protein profiles of drugs a and b, for nonzero profile norms:

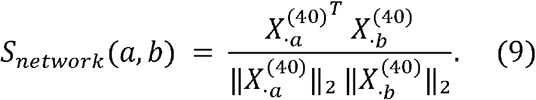

Equal-rank fusion averages the normalized ranks of M fixed scores across *N*_*pairs*_ candidate pairs. Larger scores receive higher ranks, with average ranks assigned to ties:

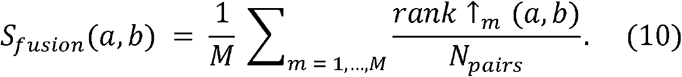

This combines fixed structural and network scores without fitting DDI annotations.^25,26^

### Two sparsification rules

The cached feature-degree cap retains single-metabolite or metabolite-pair features associated with no more than K proteins:

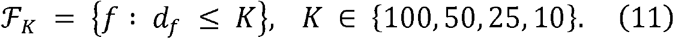

The stricter reconstruction first computes the union of predicted protein targets for each individual metabolite m across all features containing that metabolite:

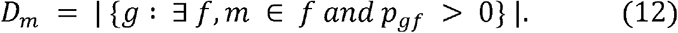

A feature is retained only when every constituent metabolite has individual degree below 100:

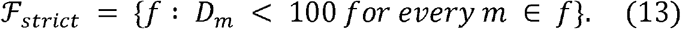

Thus, a pair feature (m,n) survives only if both *D*_*m*_ < 100 and *D*_*n*_ < 100. This differs from imposing a cap on the degree of the pair feature itself. In the supplied background, 149 of 305 observed metabolites passed the individual-metabolite rule, leaving 330 retained features and 2,956 active proteins, including 2,954 proteins in the largest connected component. These counts characterize the available reconstruction rather than a complete human interactome.

### Literature audit and reproducibility

Using ChatGPT 5.6 sol, searched the literature for 497 distinct pairs in the confidence-weighted top list that were absent from both frozen DDInter and KEGG. Two pair-specific searches per candidate were supplemented by synonym and targeted follow-up searches. Abstracts, indexed passages, and available primary full texts were classified as measured human interactions, suspected human interactions, preclinical interactions, tested combinations without a resolved interaction conclusion, negative evidence, or other evidence. This candidate-wide search was not an exhaustive systematic review. Cell-model synergy was not treated as a clinical DDI label, and an endpoint-specific negative result was not treated as proof of universal noninteraction. Reference labels were unchanged. Twenty-three candidates were present in the expanded saved DDInter snapshot; this is a reference-coverage sensitivity rather than prospective validation. Tables S5–S8 provide candidate classifications, evidence sources, search provenance, and the expanded-reference reconciliation.

## Results

### Principal findings from the complete cohort and protein subsets

The structural rankings enriched documented DDIs in the complete cohort and several protein-defined subsets (Table 1). Full-cohort top-5% precision was 49.31% against frozen DDInter, compared with 29.60% background. In the shared-kinase population, COLIG ranking reached 56.10% versus 37.35% background. Among pairs containing at least one FDA-listed P450 substrate, precision reached 62.84% against expanded DDInter versus 45.65% background.^35^ Each subset is compared with its own eligible population. The expanded DDInter reference contained 8,555 positives among 21,945 pairs (38.98%); top-5% precision was 51.58%. KEGG precision was 25.06% versus 14.67% background. These values measure recovery of the specified reference annotations, not uniformly confirmed clinical interactions.

**Table 1.**
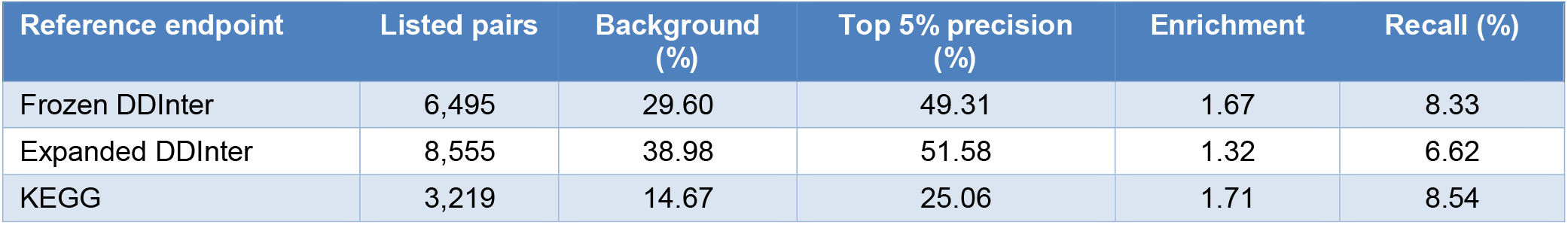
Current confidence-weighted local ranking over all 21,945 pairs.

| Reference endpoint | Listed pairs | Background (%) | Top 5% precision (%) | Enrichment | Recall (%) |
|---|---|---|---|---|---|
| Frozen DDInter | 6,495 | 29.60 | 49.31 | 1.67 | 8.33 |
| Expanded DDInter | 8,555 | 38.98 | 51.58 | 1.32 | 6.62 |
| KEGG | 3,219 | 14.67 | 25.06 | 1.71 | 8.54 |

**Figure 2.**
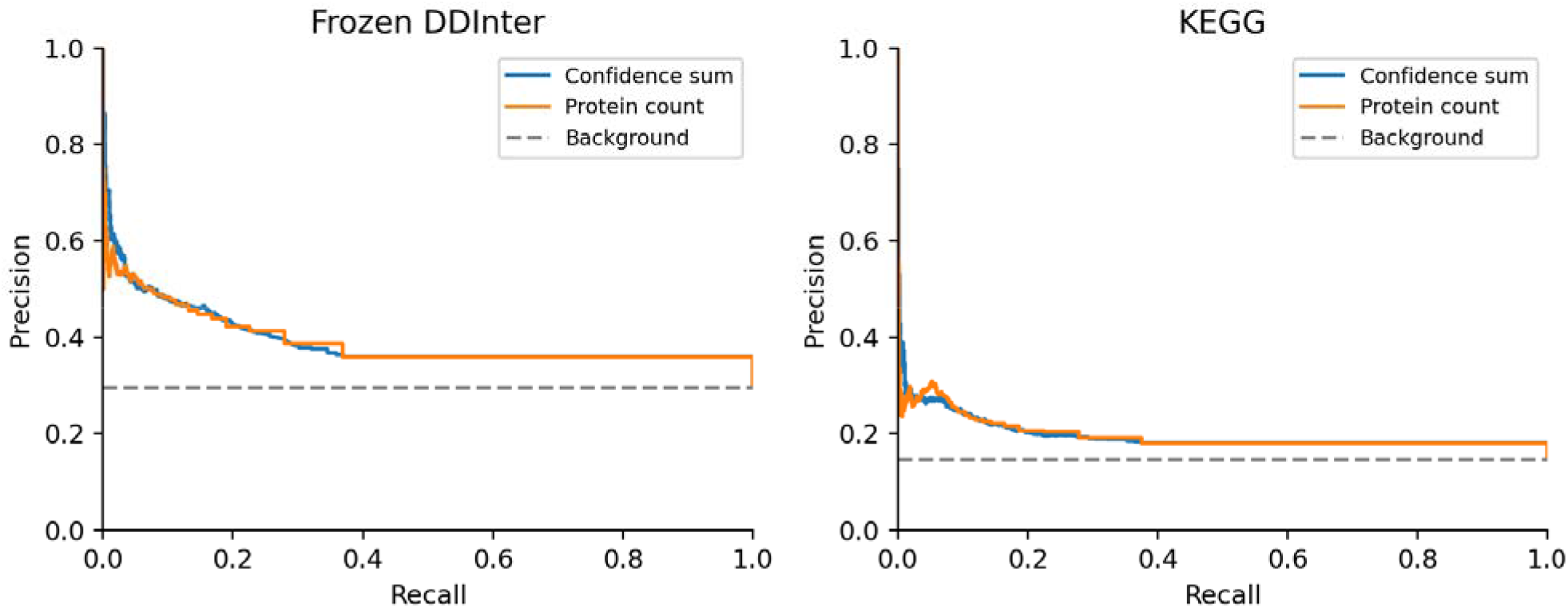
shows the precision–recall curves for the frozen DDInter and KEGG references. Precision–recall curves for fixed confidence-sum and distinct-protein-count scores. Dashed lines show prevalence in each complete pair universe. Curves include tied scores as groups and do not reorder ties using labels. The primary comparison is the top 5% of ranked drugs reported in Table 1.

### Comparison of homology filtered prediction releases

The original prediction release, which excluded templates above 35% target–template sequence identity, contained 4,083 supported drug pairs, including 1,581 frozen-DDInter and 805 KEGG positives. The current release applies a 30% cutoff. Both releases enriched documented interactions (Table 2), but they differ in coverage and aggregation as well as sequence-identity filtering. This comparison is therefore a historical sensitivity analysis, not a controlled test of the cutoff alone.

**Table 2.** Prediction releases and historical comparisons.

| **Sequence identity cutoff** | **DDInter top 5% precision (%)** | **KEGG top 5% precision (%)** |
|---|---|---|
| 30% | 49.31 | 25.06 |
| 35% | 48.11 | 27.39 |

### Mechanism specific differences in interaction enrichment

The frozen DDInter reference contained 2,306 pharmacokinetic pairs, 2,206 potentiation-tagged pairs, 231 antagonism-tagged pairs, and 1,839 pairs with unspecified mechanism. A drug pair can carry more than one mechanism flag. For each category, we compared its positives with the common set of 13,390 pairs unlisted in both DDInter snapshots. The resulting candidate populations— not positive counts—are reported in Table 3. Local top-5% precision was 32.24% versus 14.69% background for pharmacokinetics and 26.67% versus 14.14% for potentiation. The corresponding excess-precision 95% intervals were +6.52 to +29.08 and +3.85 to +23.48 percentage points. Antagonism and unspecified mechanisms had intervals including zero. Expanded-positive sensitivities retained this qualitative distinction but reduced enrichment to 1.90 for pharmacokinetics and 1.45 for potentiation.

**Table 3.** Mechanism-specific local ranking against unlisted pairs^a^.

| Mechanism category | Candidate pairs | Background (%) | Top 5% precision (%) | Fold enrichment | Excess precision 95% interval (%) |
|---|---|---|---|---|---|
| Pharmacokinetic | 15,696 | 14.69 | 32.24 | 2.19 | 6.52 to 29.08 |
| Potentiation | 15,596 | 14.14 | 26.67 | 1.89 | 3.85 to 23.48 |
| Antagonism | 13,621 | 1.70 | 2.35 | 1.39 | −1.34 to 4.02 |
| Unspecified | 15,229 | 12.08 | 14.11 | 1.17 | −4.93 to 10.14 |

<sup>a</sup> Intervals describe excess precision, the selected reference-positive fraction minus the background fraction, in percentage points. Chemical-group bootstrap resampling accounts for repeated drugs. Potentiation is a mechanism annotation, not an assay of anticancer synergy or benefit.

### P450 substrate containing pairs yield higher observed screening precision

Among the 5,474 pairs containing at least one of the 28 FDA-listed P450-substrate drugs, local top-5% precision was 62.84% against expanded DDInter versus 45.65% background (Table 4).^35^ Frozen-DDInter precision was 58.46% versus 32.70% background. This subset therefore had higher observed shortlist precision than the complete cohort, whose frozen-DDInter precision was 49.31%, but its reference prevalence was also higher. These are shortlist precisions, not whole-cohort classification accuracies.

**Table 4.** Top 5% precision in pairs containing a documented P450 substrate.

| Type of score | Candidate pairs | Background (%) | Top 5% precision (%) |
|---|---|---|---|
| Expanded DDInter local score | 5,474 | 45.65 | 62.84 |
| Expanded DDInter local plus P450 drug–drug support | 5,474 | 45.65 | 64.30 |
| Expanded DDInter local plus P450 drug–metabolite support | 5,474 | 45.65 | 62.84 |
| Frozen DDInter local score | 5,474 | 32.70 | 58.46 |

The cohort includes every pair containing at least one of 28 drugs identified as substrates in the FDA example list. Adding P450 drug–drug support by fixed rank fusion increased expanded-DDInter top-5% precision from 62.84% to 64.30%. This favorable point change was 1.46 percentage points, with a paired 95% interval of −8.15 to +13.81; it did not establish an incremental improvement. Adding shared drug–metabolite support left precision at 62.84%. Removing all P450 contributions gave 64.93%, also without an established difference. The robust descriptive finding is useful approximately 60% precision in the substrate-containing population; the present feature additions do not identify a uniquely predictive P450 contribution.

The core-P450 drug–drug COLIG union supported 145 of these pairs: 85 had expanded DDInter annotations, 40 had a metabolism flag, and 16 had a potentiation flag. The metabolism fraction was 27.59% versus 19.16% in the substrate-containing population. Its group-bootstrap excess interval, −7.44 to +25.32 points, included zero despite a nominal uniform-pair P value of 0.0079. Attractive and repulsive support selected 34 and 134 pairs, respectively, and neither sign preferentially recovered the potentiation flag. The shared drug–metabolite core-P450 score supported 29 pairs. Its metabolism and potentiation fractions were 20.69% and 10.34%, compared with backgrounds of 19.16% and 14.94%.

Current mixed records covered only CYP2A6 and CYP2C19 in the ten-enzyme core panel. Strict alignment to a documented substrate enzyme left 13 pairs, all anchored by voriconazole at CYP2C19. Five were metabolism-tagged, or 38.46%, compared with 39.71% among all 209 voriconazole pairs. Direct substrate-enzyme alignment was broader, with 21 metabolism-tagged pairs among 61 structural candidates. These limited-coverage analyses suggest specific enzyme hypotheses but do not establish that the recorded clinical interactions are mediated by the nominated P450.

### Protein classes reveal numerical concentration and ranking redundancy

All 3,916 proteins in the prediction universe were assigned to primary functional classes. In populations where both drugs had at least one predicted target in the class, the complete local score achieved 59.43% precision for ion channels versus 38.58% background, 52.61% for GPCRs versus 31.81%, and 54.47% for other oxidoreductases versus 32.57%. Chaperone-, phosphatase-, and isomerase-engaging populations had point estimates near 60%, but broad intervals prevented ranking them as superior classes. The highest estimate, 74.36% for lipid-kinase engagement, came from a shortlist of only 39 weighted pairs and was unstable. Class populations overlap, and these rankings use the complete local score, including evidence from other classes (Table 5).

Protein kinases contributed 83.45% of confidence weight in the complete local top-5% list. Kinase-only ranking achieved 48.67% precision, and its top list overlapped the complete list by 80.39%. Removing all kinase contributions nevertheless retained 48.23% precision, a decrease of approximately 1.07 percentage points with a paired 95% interval of −5.46 to +3.85 points. A large numerical contribution therefore did not establish a unique predictive contribution: other protein classes ranked many of the same drug pairs highly.

### COLIG ranking advantages extend beyond kinases with class specific exceptions

At equal top-100 drug-pair budgets against frozen DDInter, confidence-ranked COLIG precision was 67.00% for kinases versus 39.88% for NoCOLIG drugs, 60.90% versus 34.47% for other hydrolases, 55.00% versus 39.43% for other transferases, 53.00% versus 36.41% for nuclear receptors, and 69.00% versus 37.00% for other proteins. Each COLIG value also exceeded its class background (Table 6). These observed differences of approximately 16–32 percentage points extend the favorable pattern beyond kinases, but do not establish independent class-specific superiority after simultaneous correction.

**Table 5.** Complete local ranking in selected protein-class populations.

| Both drugs engage class | Candidate pairs | Background (%) | Top 5% precision (%) | Excess precision 95% interval (pp) |
|---|---|---|---|---|
| Chaperones | 2,145 | 39.67 | 60.61 | -3.78 to 44.71 |
| GPCRs | 9,453 | 31.81 | 52.61 | 6.58 to 35.81 |
| Ion channels | 4,005 | 38.58 | 59.43 | 2.37 to 38.20 |
| Isomerases | 3,741 | 41.89 | 59.90 | -2.68 to 36.37 |
| Other oxidoreductases | 9,180 | 32.57 | 54.47 | 8.44 to 37.21 |
| P450 family | 2,701 | 38.69 | 46.65 | -9.27 to 30.08 |
| Phosphatases | 3,403 | 41.93 | 59.95 | -2.48 to 37.32 |
| Protein kinases | 12,246 | 34.04 | 51.49 | 3.77 to 30.39 |

Each population includes every pair among drugs predicted to bind at least one member of the class in percentage points(pp). The complete local score is used, including support from other classes. Populations overlap; their raw precisions cannot rank class specificity. All classes and class-only scores are supplied separately.

**Table 6.** Top 100 COLIG and explicit same pocket NoCOLIG comparisons.

| Protein class | Candidate pairs | NoCOLIG supported pairs | Baseline (%) | COLIG top 100 precision (%) | NoCOLIG top 100 precision (%) | Difference 95% interval (pp) |
|---|---|---|---|---|---|---|
| Protein kinases | 7,609 | 124 | 37.4 | 67.0 | 39.9 | -4.5 to 50.5 |
| Nuclear receptors | 6,010 | 125 | 32.6 | 53.0 | 36.4 | -20.5 to 53.9 |
| Other hydrolases | 4,007 | 164 | 38.6 | 60.9 | 34.5 | -4.0 to 49.8 |
| Other transferases | 4,161 | 161 | 37.6 | 55.0 | 39.4 | -13.0 to 44.6 |
| Other oxidoreductases | 3,266 | 133 | 37.7 | 32.5 | 32.4 | -25.2 to 27.6 |
| Other proteins | 7,296 | 296 | 35.2 | 69.0 | 37.0 | 1.8 to 58.7 |
| Pooled all proteins | 12,671 | 811 | 32.4 | 66.0 | 34.0 | -3.0 to 55.6 |

Frozen DDInter; equal budgets of 100 weighted drug pairs. The final column is the paired COLIG-minus-NoCOLIG precision interval in percentage points(pp). Candidate populations and backgrounds differ by class. Intervals are pointwise; no class advantage survived the corresponding simultaneous adjustment. A pair may have both assignment types on different structures.

The archives contained 131,051 COLIG and 2,536 NoCOLIG drug-pair–protein supports. In the 7,609-pair shared-kinase population, every pair selected by the complete local top-5% score had kinase COLIG support. Kinase-COLIG-only ranking gave the same 56.10% precision versus 37.35% background. The NoCOLIG control contained only 124 kinase pairs, insufficient to fill the same top-5% budget. The equal top-100 comparison therefore supplies the direct control without introducing zero-score pairs.

All six primary classes with at least 100 supported pairs for each assignment type had numerically higher confidence-ranked COLIG precision. The sixth class, other oxidoreductases, was essentially tied at 32.50% versus 32.36%, with both below its 37.66% background. Thus, five classes showed substantial point differences and one showed a near tie, rather than a uniformly useful result.

When at least 20 pairs are considered, confidence-ranked COLIG precision was higher in 16 of 17 classes against the full set of frozen DDInter, all 17 against expanded DDInter, and 16 against KEGG. Proteases were the frozen-reference exception. These concordant directions extend the observed tendency beyond the six top-100 classes, although overlapping drug pairs prevent treating classes or reference representations as independent replications. Protein-count rankings were less uniform: the frozen-DDInter hydrolase top-100 contrast changed from +26.43 percentage points with confidence weighting to −4.21 with counting. The assignment type, confidence weighting, and protein class all contribute to the descriptive pattern.

Pooling protein classes yielded 12,671 pairs sharing at least one predicted target, including 6,685 COLIG-supported and 811 NoCOLIG-supported pairs. At the primary top-5% cutoff, confidence-ranked COLIG precision was 50.98% versus 36.58% for NoCOLIG, compared with 32.44% background. The paired difference was +14.40 percentage points (95% interval −1.36 to +30.53). At top 100, precision was 66% versus 34% (difference interval −2.97 to +55.60 points). Both intervals included zero.

### Distinct protein counts and confidence both contribute to ranking

The number of distinct supporting proteins carried useful predictive information. Across COLIG-supported pairs, as shown in Table 7, the frozen-DDInter fraction rose from 29.24% for one supporting protein to 30.71% for two, 36.17% for three to five, 39.46% for six to ten, 44.02% for 11 to 25, 44.36% for 26 to 50, and 52.22% for more than 50. NoCOLIG pairs showed no comparably orderly trend, although the high-count bins were sparse. These are fractions among supported pairs, not top-list precisions or a biological dose-response measurement. The positive association suggests that repeated COLIG support across different proteins contains information beyond the presence of a single predicted assignment and may play a role in seeding responses across different parts of the cellular network

**Table 7.** Documented drug interactions by distinct supporting protein count^a^.

| Supporting proteins | COLIG pairs | COLIG listed fraction (%) | NoCOLIG pairs | NoCOLIG listed fraction (%) |
|---|---|---|---|---|
| 1 | 2,001 | 29.2 | 446 | 36.5 |
| 2 | 1,133 | 30.7 | 177 | 35.6 |
| 3-5 | 1,443 | 36.2 | 105 | 42.9 |
| 6-10 | 816 | 39.5 | 48 | 25.0 |
| 11-25 | 527 | 44.0 | 23 | 52.2 |
| 26-50 | 133 | 44.4 | 6 | 16.7 |
| 51+ | 632 | 52.2 | 6 | 16.7 |

<sup>a</sup> Frozen DDInter; all positively supported pairs, without shortlist selection. Each assignment type counts its distinct supporting human proteins across structural contexts. A pair can appear in both assignment types on different structures or contexts. Fractions describe association within each count bin, rather than the top-5% precision, causal dose response, or count-matched estimates. The high-count NoCOLIG groups are sparse.

Confidence-based selection ranks drug pairs by the sum of retained COLIG confidences across distinct proteins, counting each protein once. The highest-scoring 5% are selected, with fractional weighting of ties at the cutoff. Labels are not used in selection, and the score is not a calibrated clinical interaction probability. We also divided the sum by supporting-protein count to examine mean confidence (Table 8). In the shared-kinase population, mean-confidence ranking achieved 73.00% top-100 precision versus 39.33% for NoCOLIG. At top 5%, kinase COLIG precision was 54.21% using protein count, 56.10% using summed confidence, and 57.30% using mean confidence, all above 37.35% background. The mean-versus-sum difference of +1.20 percentage points had a 95% interval of −9.18 to +12.82, so it did not establish a superior predictor. At a smaller equal number of cases considered, mean-confidence COLIG precision exceeded NoCOLIG in 16 of 17 classes against frozen DDInter, all 17 against expanded DDInter, and 16 against KEGG; no class contrast survived the corresponding simultaneous adjustment. Top-100 results are secondary and should not replace the primary top-5% comparison.

**Table 8.** Separating supporting protein count from confidence based ranking ^a^.

| Population | Ranking score | COLIG top 5% precision (%) | COLIG top 100 precision (%) | NoCOLIG top 100 precision (%) |
|---|---|---|---|---|
| Pooled shared proteins | Confidence sum | 51.0 | 66.0 | 34.0 |
| Pooled shared proteins | Protein count | 52.1 | 54.7 | 35.3 |
| Pooled shared proteins | Mean confidence | 48.9 | 59.5 | 35.5 |
| Shared kinase | Confidence sum | 56.1 | 67.0 | 39.9 |
| Shared kinase | Protein count | 54.2 | 52.0 | 39.6 |
| Shared kinase | Mean confidence | 57.3 | 73.0 | 39.3 |

<sup>a</sup> Frozen DDInter. Pooled shared-protein population: 12,671 pairs, 32.44% background. Shared-kinase population: 7,609 pairs, 37.35% background. Top 5% and top 100 are involve different number of proteins. NoCOLIG can fill the top-100 budget in both populations but cannot fill the kinase top-5% case cutoff, so that padded metric is omitted. Mean confidence divides the sum by the number of distinct supporting proteins. These are fixed exploratory scores, not trained or independently optimized predictors. Point advantages do not establish superiority: the kinase mean-versus-sum top-5% difference interval includes zero.

Protein count and summed confidence were strongly related among positively supported pairs: Spearman correlations were 0.906 for COLIG and 0.951 for NoCOLIG. COLIG pairs averaged 19.60 distinct supporting proteins (median 3), compared with 3.13 (median 1) for NoCOLIG; kinase means were 26.13 and 2.56, again with medians 3 and 1. Count–DDI correlations were more modest, 0.141 and 0.001, respectively. Table 7 reports the count-stratified fractions and sample sizes. Together, these analyses show contributions from both count and confidence without establishing an independent assignment-type effect.

### Shared top pocket binding and direct COLIG evidence

We asked whether predicting a direct COLIG interaction between two drugs provides more information about clinical drug–drug interactions than simply predicting that both drugs bind to a protein’s top-ranked pocket. For each protein structure, we formed all drug pairs from its individual binding predictions, yielding 8,623 pairs across the available structures. Of these, 6,685 had explicit COLIG support in at least one structure, while 811 had explicit NoCOLIG support; these groups can overlap when assignments differ between structures. We ranked all methods within the same 8,623-pair population, using the same confidence rule to make the comparison fair. Among the highest-ranked 5% of pairs, documented DDInter interactions accounted for a precision of 52.42% using shared top-pocket binding, 54.04% using COLIG support, and 36.10% using explicit NoCOLIG support, compared with 34.54% in the full comparison population. Thus, COLIG-supported pairs had higher observed precision, particularly relative to NoCOLIG pairs, but the uncertainty intervals did not establish a reliable advantage. These results suggest that shared top-pocket binding already contains useful predictive information.

### Noninterface pockets supply most of the complete ranking signal

Noninterface single-chain COLIGs within dimers achieved 50.15% full-cohort precision, isolated PDB monomers 49.01%, and AlphaFold2 monomers 45.43%, compared with a 29.60% background precision (Table 9). Drugs predicted to bind outside the dimer interface and isolated-monomer top lists overlapped by 84.19%, and the dimer-chain list overlapped the full list by 94.27%. Of the full top-list confidence weight, 58.03% was assigned to noninterface dimer chains, 32.89% to isolated PDB monomers, 8.90% to AlphaFold2 monomers, and 0.18% to interface-adjacent dimer pockets (Figure 3). These shares depend on coverage, confidence, and maximum-per-protein allocation; they do not measure biological importance. Attractive-only confidence achieved 39.88% precision with 1,902 supported pairs, repulsive-only confidence 49.24% with 6,391, and their union 49.31%. Unequal coverage and correlated evidence prevent interpreting the small union increment as proof that attractive interactions are biologically irrelevant.

**Table 9.** Structural context of COLIG evidence^a^.

| Structural context<br>COLIG pairs |  | NoCOLIG pairs | Full cohort top 5% precision (%) | Within context precision / background (%) | Top list confidence contribution (%) |
|---|---|---|---|---|---|
| Isolated PDB monomer | 3,663 | 235 | 49.01 | 58.21 / 38.16 | 32.89 |
| AF2 monomer | 1,529 | 128 | 45.43 | 51.63 / 43.59 | 8.90 |
| Non-interface chain in dimer | 5,877 | 636 | 50.15 | 56.31 / 35.07 | 58.03 |
| Interface-adjacent dimer | 494 | 15 | Insufficient support | 76.81 / 52.08 | 0.18 |
| All contexts | 6,685 | 811 | 49.31 | 53.58 / 34.54 | 100.00 |

<sup>a</sup> Frozen DDInter. Full-cohort comparisons rank all 21,945 pairs against 29.60% background. Within-context comparisons rank the top 5% of that context’s pooled top-pocket pair population; their budgets and backgrounds differ. Interface-adjacent native COLIG precision is 76.81% (95% interval 45.19–100.00%) at only 27.65 selected pair mass among 553 candidates, versus 52.08% background; its excess interval includes zero. Only 494 interface-adjacent COLIG pairs and 15 explicit NoCOLIG pairs are supported. Confidence contribution allocates each pair/protein maximum to its supporting context, splitting exact ties; it measures score composition, not causation. Units mapping to one human protein and a 247-common-protein comparison are separate sensitivities.

**Figure 3.**
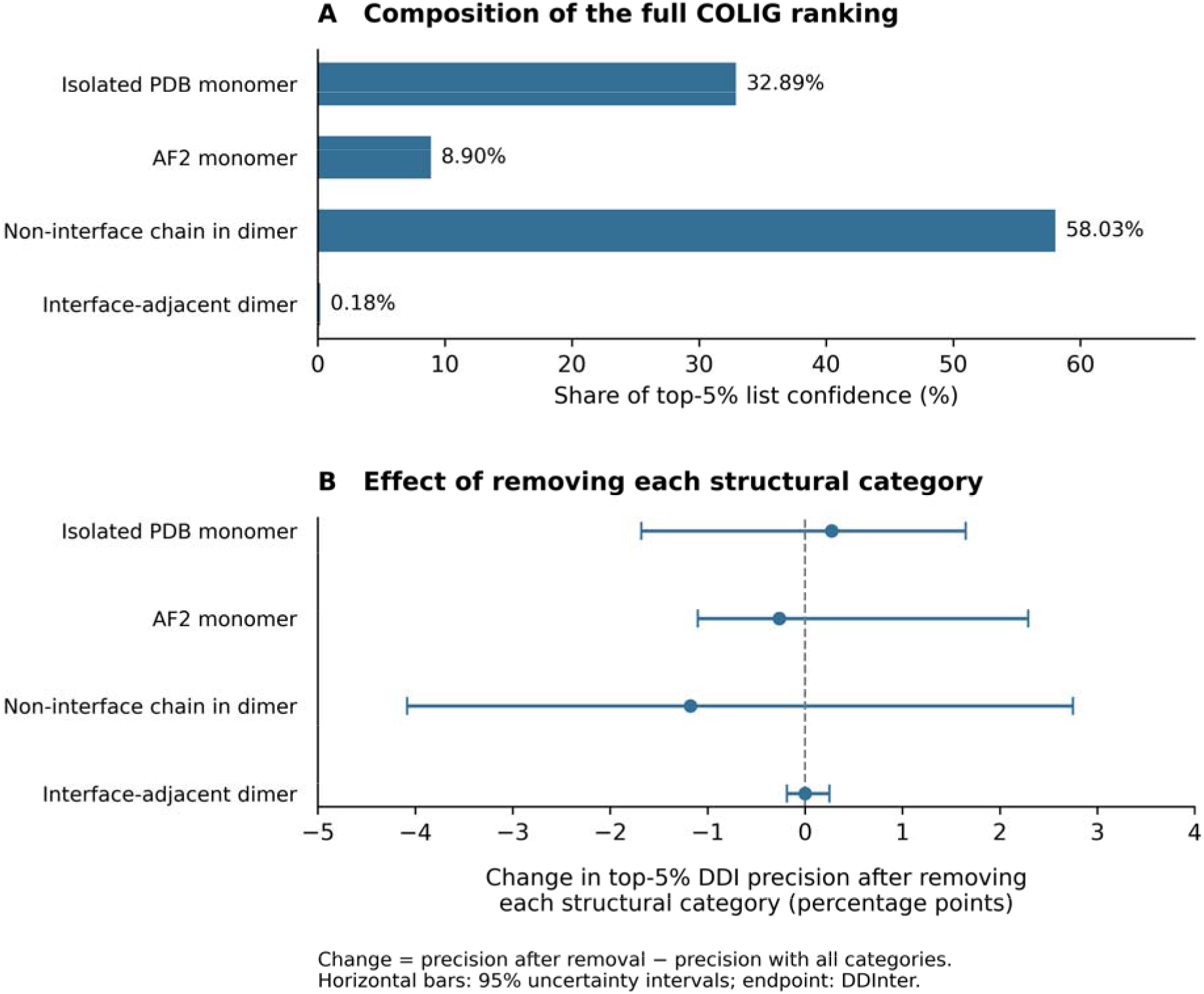
Structural-context composition and ablation of the complete COLIG ranking. (A) share of native-confidence weight in the full-cohort top 5%, assigning each pair/protein maximum to its supporting context and dividing ties equally. (B) change in frozen-DDInter top-5% precision after deleting each context; bars show 1,000 paired chemical-group bootstrap 95% intervals. All intervals include zero. Contribution depends on coverage and confidence and does not establish causal importance. Interface-adjacent pockets have a high but uncertain within-subset precision estimate (Table 9), despite their small contribution to the complete ranking.

The subset of binding drugs adjacent to the protein-protein interface contained 553 pooled candidate pairs, including 494 COLIG-supported pairs and only 15 explicit NoCOLIG pairs. For the top 5% of this population, supplied COLIG-confidence ranking achieved 76.81% precision versus 52.08% background, selecting 27.65 weighted pairs. Its precision interval was broad (45.19–100.00%), and the excess-precision interval included zero. This sparse subset cannot fill the full-cohort budget or support a stable NoCOLIG top-100 comparison. Its high point estimate is biologically interesting in light of the enrichment of disease-associated variants in interfacial ligand-binding pockets reported by Gao, Zhou, and Skolnick, but the two observations do not establish a shared mechanism.^39^

### Drug deletion tests and Liverpool interaction assessments

Removing each drug in turn retained supplied-confidence COLIG top-5% precision between 46.91% and 50.70%, always above the corresponding DDInter background. Removing dasat nib produced the largest decrease, whereas removing ethanol produced the largest increase. Deleting each of the 209 chemical groups gave the same range. Shared top-pocket binding precision ranged from 48.77% to 52.15%. Thus, enrichment did not depend entirely on one drug. These deletion ranges are not confidence intervals and do not establish performance on additional drugs.

We also evaluated the rankings against Liverpool interaction assessments, using 1,566 drug pairs for which the available assessments did not conflict. Of these pairs, 46.42% carried an interaction warning. Among the highest-ranked 5%, the warning rate was 80.84% for COLIG predictions ranked by their supplied pair-confidence scores, 75.73% for shared top-pocket binding drugs, and 77.01% for COLIG predictions scored using the same individual-drug confidence rule as the shared-binding comparison. The supplied COLIG scores exceeded shared binding by 5.11 percentage points, but the 95% interval for this difference ranged from −10.50 to +16.07 points, so an advantage was not established. Excluding yellow assessments and comparing only red or amber warnings with green assessments left 1,460 pairs; COLIG precision remained 79.45%, compared with a warning rate of 42.53% in that population. These assessments involve a limited set of primary drugs from the Liverpool COVID, HIV, and hepatitis resources, and some warnings reflect expected pharmacological risks rather than measured interactions. The results therefore show agreement with another reference resource for the same drugs, rather than validation on new drugs.

### Independent NoCOLIG protein assignments

We constructed an independent protein profile for each drug from its singleton NoCOLIG records, retaining the maximum confidence per drug–protein pair across structures. Pair scoring then required shared proteins, not occurrence in the same pocket or structural unit. Profiles covered 182 of 210 drugs, with 7,468 assignments to 3,461 proteins; 4,382 drug pairs shared at least one assigned protein. Shared-protein count achieved 41.03% frozen-DDInter top-5% precision versus 29.60% background, or 1.39-fold enrichment (excess-precision interval +0.53 to +21.25 percentage points). Confidence-weighted overlap gave 40.70% precision (excess interval +0.22 to +21.15), without improving the count result. For expanded DDInter, count precision was 46.77% versus 38.98% background, with an excess interval including zero; KEGG precision was 13.57% versus 14.67%, showing no enrichment. Independent NoCOLIG target overlap therefore captures part of the frozen-DDInter association, with a lower point precision than the 49.31% direct COLIG ranking and no established causal interpretation.

### Network propagation from independent NoCOLIG seeds

To test whether network context improved prediction from independent NoCOLIG target assignments, we used each drug’s maximum-confidence NoCOLIG protein profile as the seed for the same STRING physical-interaction, feature-degree-cap-100 metabolite, and combined protein networks examined above. Seed profiles were normalized to unit mass, propagated for 40 iterations with a restart probability of 0.35, and drug pairs were ranked by cosine similarity between their propagated profiles. Against frozen DDInter, the unpropagated NoCOLIG shared-confidence score gave 40.70% for the top-5% precision, whereas STRING, metabolite, and combined-network propagation gave 36.36%, 37.75%, and 36.82%, respectively. Equal-percentile-rank fusion of the local score with the metabolite-network score gave the best network-containing result, 40.01%, a change of −0.69 percentage points relative to the local score, with a paired 95% interval of −3.85 to +1.61 points. For expanded DDInter, metabolite-network fusion increased precision from 46.69% to 47.21%, but the +0.52-point difference had an interval of −2.62 to +2.91 points. STRING propagation increased the KEGG point estimate from 13.51% to 15.58%, but its +2.08-point difference also remained unresolved (−2.12 to +7.36 points). Thus, propagating the independent NoCOLIG assignments did not establish incremental predictive value over direct protein overlap and potentially diluted the frozen-DDInter signal.

### COLIG protein assignments and network propagation

We also represented each drug by proteins supported by at least one COLIG record, forming unsigned, attractive-only, and repulsive-only profiles. Ranking pairs by shared-protein count gave frozen-DDInter top-5% precisions of 51.11%, 51.26%, and 51.28%, respectively, versus 29.60% background. Corresponding precisions were 52.56%, 52.53%, and 52.69% for expanded DDInter and 29.89%, 30.98%, and 29.98% for KEGG. Thus, shared COLIG-derived protein assignments contain a substantial ranking signal, with little difference between the structural signs.

We then propagated the three COLIG protein profiles through the STRING physical-interaction network and the STRING combined with the feature-degree-cap-100 metabolite network. For frozen DDInter, STRING precision ranged from 47.41% to 48.85%, and STRING plus metabolites ranged from 48.12% to 49.40%; both were below the corresponding COLIG-only scores. Propagation modestly increased the expanded-DDInter point estimates, with STRING giving 54.34–54.86% precision and STRING plus metabolites giving 53.70–54.23%, but every paired 95% interval for the change from the local score included zero. KEGG likewise showed no resolved improvement: the attractive STRING-plus-metabolite score was essentially unchanged at 30.99% versus 30.98% locally, and the other propagated scores were lower. Overall, the predictive association is concentrated in direct shared COLIG protein assignments. Network propagation broadens this signal but does not provide statistically resolved incremental predictive value under the present analysis.

### Literature review shows why unlisted pairs cannot be treated as confirmed negatives

Among the 497 candidates absent from frozen DDInter and KEGG, the audit identified four pairs with measured human interaction evidence: gefitinib–sorafenib, imatinib–sorafenib, nelfinavir– saquinavir, and caffeine–clopidogrel. Sorafenib reduced gefitinib exposure in a phase I trial; imatinib exposure with sorafenib was higher than historical single-agent values; nelfinavir increased saquinavir exposure; and caffeine altered selected platelet-inhibition measurements after clopidogrel.^40,41,42,43^ A colchicine–sunitinib case report provided suspected human evidence, with renal impairment and high colchicine exposure limiting causal attribution.^44^ The audit also found 37 preclinical interaction candidates, 33 tested combinations without a resolved interaction conclusion, and 413 pairs without qualifying empirical evidence located (Tables S5–S7). Other categories included endpoint-specific negative evidence, different chemical forms, and computational hypotheses; evidence could overlap. These categories cannot be pooled into a revised clinical precision estimate. Twenty-three candidates were present in expanded DDInter (Table S8), illustrating that frozen-reference nonlisting is not a biological negative.

We then asked whether the structural predictions could distinguish highly ranked pairs with documented interactions from highly ranked pairs absent from the references. The two groups had similar prediction-confidence scores and similar patterns of predicted involvement in biological pathways. Comparing pairs that shared one drug and had similar local ranking scores also failed to identify a consistent distinguishing feature among those examined. Excluding drugs predicted to bind more than 100 proteins did not consistently improve performance either. These results suggest that incomplete reference coverage may contribute to the apparent prediction errors, and that the additional pathway features may largely repeat information already captured by the local score. However, they do not establish that every unlisted prediction is a real interaction. They also leave open the possibility that important biological interactions are missing from the features tested.

## Conclusions

Predicted top-pocket engagement provides a useful preliminary screen for documented drug– drug interactions without fitting clinical labels or experimentally curated drug-target profiles. Pooled target-overlap proxies retain the top-pocket restriction of the supplied predictions and also enrich documented DDIs. Direct attractive or repulsive COLIG scoring is not uniformly superior to those proxies in the complete cohort. The positive result is stronger than a pooled drug pair average alone convey shared-kinase COLIGs achieved 56.1% top-5% precision versus 37.4% background, and the P450-substrate-containing population achieved 62.8% against expanded DDInter versus 45.7% background. At equal top-100 considered pairs of proteins , confidence-ranked COLIG precision substantially exceeded explicit same-pocket NoCOLIG precision in five of six evaluable classes, with oxidoreductases essentially tied. The favorable direction extended to 16 of 17 classes. Mean-confidence kinase ranking retained the advantage, achieving 57.3% for the top 5% and 73.0% for the top 100, while protein-count stratification showed increasing documented-DDI fractions with broader COLIG support.

Structural-unit matching and jackknife analyses retain the pocket-engagement association: isolated PDB monomers and non-interface dimer-chain pockets achieved 49.0% and 50.1% complete-cohort top-5% precision, respectively, versus 29.6% background. Interface-adjacent pockets supplied only 0.18% of top-list confidence, and their removal left overall precision unchanged. Their high within-subset estimate, 76.8% versus 52.1% background, remains an interesting but sparse and uncertain result. These observations support pocket engagement across structural association contexts without establishing that direct protein–protein-interface perturbation drives the overall enrichment. Direct COLIG interactions and noncontact pocket engagement are candidate functional routes; conformational change and altered protein–protein association have not been demonstrated. These findings motivate a qualitative picture in which local pocket relationships, accumulated across distinct proteins, help identify drug pairs capable of producing a physiological interaction. The tested explicit network and pathway additions provided no consistent incremental benefit, suggesting that structural engagement profiles already summarize useful distributed information for this task. This does not establish that cellular networks are unimportant downstream. The more specific comparison distinguishes attractive or repulsive COLIG relationships from explicit same-pocket NoCOLIG assignments. Its favorable class-level point estimates motivate additional contributions from direct ligand–ligand relationships, but do not establish a uniform independent advantage. Class-dependent effects of direct COLIG interaction and noncontact pocket engagement remain plausible. Sparse controls, overlapping pairs, exploratory score comparisons, and inconclusive count-matched results require caution about causation and independent class superiority. Independent validation and targeted perturbation experiments are needed to determine which nominated pocket relationships actually initiate the recorded interactions.

### Local observations can encode distributed biology

The term local applies to the physical source of each predicted observation. It does not mean that the drug-pair score describes one pocket, one protein, or a cell-independent physiological outcome. Aggregation across many targets can summarize distributed engagement, including pathway relationships, without an explicit random walk. The class-ablation and convergence results make this distinction concrete: heavily weighted classes are not necessarily uniquely informative, and small-weight classes can preserve much of the pair ranking. It is physically reasonable that an initiating pocket event could alter competition, conformation, transport, or catalytic turnover and thereby propagate to physiology. That possibility motivates the method. The present evidence supports the existence of a useful structural association, not the proposition that most DDIs begin at the nominated pocket or that network processes contribute little to the eventual clinical effect. The similarity of predictions from several representations is compatible with shared biology, target burden, correlated prediction error, and common annotation bias. These explanations require different validation experiments.

### Functional responses may depend on the protein class

The class comparisons suggest a useful mechanistic hypothesis: direct COLIG interaction and noncontact engagement of the same pocket may differ in their ability to perturb particular protein functions. The observed top-100 advantage of kinase COLIGs, 67.0% versus 39.9% for NoCOLIG, is consistent with a role for the joint structural relationship beyond merely assigning both drugs to a pocket. Substantial advantages in four additional classes motivate testing how this relationship depends on protein function, pocket geometry, and ligand chemistry. The essentially tied oxidoreductase result identifies an informative exception rather than negating the other observations.

Both mechanisms could operate. Direct attractive or repulsive relationships may alter co-occupancy, ligand positioning, or protein conformation. Drugs assigned to the same pocket without direct contact may also modify function through occupancy or conformational changes. The present assignment files do not by themselves establish simultaneous occupancy, competition, inhibition, or activation. In particular, predicted kinase binding should not automatically be called kinase inhibition. The class patterns provide targets for functional tests; they do not identify the downstream physiological consequence of any one drug pair.

The count and confidence analyses sharpen this interpretation. The rising DDI fraction with distinct COLIG-supporting proteins suggests that distributed engagement can matter even when each contributing observation is local. The 73.0% kinase top-100 precision after averaging confidence shows that useful ranking persists after removing the explicit count multiplier. However, mean confidence can remain associated with coverage and annotation, and the exact count-matched comparisons did not establish an independent assignment-type effect across all supported pairs. Differences in structural coverage, confidence distributions, drug composition, and clinical annotation remain alternatives to class-dependent biology. These alternatives delimit the proposed mechanism without obscuring the positive screening result. The current stratification concerns protein target classes; ligand chemical and therapeutic classes require a separate analysis.

### Mechanism and protein coverage constrain interpretation

The stronger pharmacokinetic and potentiation associations show that the recorded endpoint is context dependent. A change in AUC or clearance is not the same measurement as respiratory depression, loss of seizure control, or arrhythmia risk. A CYP substrate–inhibitor relationship can increase exposure, whereas inhibition of prodrug activation can decrease an active metabolite; a structural attractive/repulsive sign does not resolve that direction. P450-class binding also includes enzymes involved in endogenous metabolism, which cannot all be treated as drug-clearance targets.

The substrate analysis illustrates why a high subgroup precision and an informative mechanism are different claims. The FDA-confirmed subset had higher precision, but a different background. The mixed-COLIG enzyme alignment reduced to one substrate anchor, and the apparent enrichment disappeared against that drug’s own pair population. Similar care is necessary for protein-class strata. High prediction coverage and extensive clinical annotation can make a class appear useful without establishing that the class mediates the DDI.

The mechanistic interpretation therefore has two levels. The observed result is that top-pocket engagement, accumulated across proteins, enriches documented DDIs in both isolated and complex-associated structural contexts. A candidate explanation is that direct attractive or repulsive COLIG relationships alter local occupancy or function, while drugs assigned to the same pocket without a direct interaction may influence function through noncontact occupancy or conformational changes. Both routes could contribute to different protein classes. The current comparisons do not establish either route for an individual drug pair, distinguish allosteric from other functional effects, or demonstrate altered protein–protein association. In particular, the larger numerical contribution of non-interface dimer-chain evidence reflects the available score support and cannot itself be interpreted as stronger collective regulation.

### Main limitations and the tests needed to distinguish explanations

The first limitation is endpoint ascertainment. Positive records mix direct pair evidence, pharmacological warnings, and class-level extrapolation, whereas unreported pairs have no uniform experimental meaning. Source reconciliation changed the baseline substantially. A source-frozen external panel with explicit evidence provenance and measured noninteractions, where available, would be more informative than repeated reanalysis of the same reference pairs.

Second, structural predictions are not occupancy or turnover measurements. Template reuse, multiple mapped chains, protein-family enrichment, and missing pose information diminish correlated evidence. The current 30%-filtered drug–drug release and earlier 35% release differ in coverage and aggregation; they are not a controlled cutoff sweep. Homology filtering reduces one source of information overlap without eliminating all bias.

Finally, the present data do not establish causation. A discriminating experimental design would perturb a nominated pocket or enzyme while preserving unrelated drug exposure, quantify binding or turnover for each agent and their combination, and test whether the expected interaction disappears with that perturbation. Protein-class or pathway association alone cannot replace this intervention. Computational counterfactuals can prioritize such experiments, but their validity depends on experimentally defensible binding, exposure, and functional assumptions.

### Future work

Individual-drug protein-engagement predictions can contribute to a preliminary pair screen, with explicit COLIG assignments adding a structural description of the candidate relationship. Where joint structural analysis is required, that additional pair information should be acknowledged; single-drug predictions alone do not establish an explicit joint pose. At the top-5% ranking most reference positives remain outside the shortlist. Unlisted predictions require follow-up because database absence can represent either missing evidence or a true failure of the prediction.

### Supporting Information

Supplementary Tables S1–S8: cohort drug identities; frozen DDInter and KEGG labels; expanded DDInter labels and mechanism flags; candidate-wide literature classifications, evidence details, search provenance, and expanded-reference reconciliation (TSV and JSON files in ZIP). Additional supplementary files contain complete protein-class comparisons, matched structural-unit and common-confidence controls, count and mean-confidence sensitivities, uncertainty and zero-padding audits, molecular-association contexts and ablations, mechanism-specific and P450 analyses, deletion tests, Liverpool assessments, and independent NoCOLIG and COLIG propagation results (TSV, JSON, and TXT files in ZIP). The separate data and code archive contains inputs and programs for reproducing the reported analyses (ZIP).

## Data and Software Availability

The supplementary materials provide the drug dictionary, frozen and expanded reference labels, mechanism flags, literature classifications and primary-source links, score tables, and provenance for the reported comparisons. The separate reproducibility archive contains structural inputs, external-reference snapshots, analysis programs, and input-path manifests, including the independent NoCOLIG and COLIG propagation analyses. Zenodo deposition: https://zenodo.org/uploads/23734029.

## Acknowledgments

This research was supported in part by a gift from the Parker H. Petit AI-Driven Drug Discovery Initiative and grant GM-118039 from the National Institute of General Medical Sciences of the National Institutes of Health. We thank Jessica Forness for reviewing the manuscript and Bartosz Ilkowski for computational support.

## Notes

The authors declare no competing financial interest.

